# Importance of Molecular Dynamics Equilibrium Protocol on Protein-lipid Interactions near Channel Pore

**DOI:** 10.1101/2021.11.28.470286

**Authors:** Wenjuan Jiang, Jerome Lacroix, Yun Lyna Luo

## Abstract

Multiscale molecular dynamics (MD) simulations using Martini coarse-grained (CG) and all-atom (AA) forcer fields are commonly used in membrane protein studies. In particular, reverse-mapping an equilibrated CG model to an AA model offers an efficient way for preparing large membrane protein systems with complex protein shapes and lipid compositions. Here, we report that this hybrid CG-equilibrium-AA-production protocol may artificially increase lipid density and decrease hydration in ion channel pores walled with transmembrane gaps. To understand the origin of this conundrum, we conducted replicas of CG, AA, and reverse-mapped AA simulations of the pore domain of the mechanosensitive Piezo1 channel in a non-conducting conformation. Lipid/water density analysis and free energy calculations reveal that the lack of initial pore hydration allows adjacent lipids to enter the pore lumen through gaps between pore helices during CG simulation. Due to the mismatch between CG and AA lipid kinetics, these pore lipids remain trapped in the subsequent AA simulations, despite unfavorable binding free energy. We tested several CG equilibrium protocols and found that a protocol restraining the whole lipid produces pore hydration consistent with AA results, thus eliminating this artifact for further studies of lipid gating and protein-lipid interactions.

**WHY IT MATTERS:** Membrane-embedded proteins constantly interact with lipid molecules. Computational molecular dynamics simulations have become an indispensable tool for investigating the role of such protein-lipid interactions. Using mechanosensitive Piezo1 channel as model, we found that subtle differences in solvation and equilibrium protocols between coarse-grained and all-atom MD simulations can result in different lipid densities inside the channel pore. We identify the underlying cause of this discrepancy and propose alternative protocols to avoid this artifact.

## INTRODUCTION

Computational molecular dynamics (MD) simulation has become an indispensable tool for ion channel research. Thanks to advances in X-ray crystallography, cryogenic electron microscopy (cryo-EM), and artificial intelligence-driven structure prediction algorithms, the number of membrane protein structures has largely increased over the past few years (1). MD simulations are often needed to study how membrane protein structures, usually obtained in non-physiological conditions, behave in the physiological environment of hydrated membranes at body temperature. Various MD simulation engines are publicly available and each of them comes with its own tools for solvation, membrane embedding, and a set of standard equilibrium protocols. In addition, multiscale MD simulations using both coarse-grained (CG) and all-atom (AA) have become popular (2). Among various CG models, Martini force field is routinely used in simulating lipid distribution and protein-lipid binding (3, 4). While enabling simulations speeds about an order of magnitude higher than AA simulations, current Martini protein models require tertiary structure constraints, hampering unbiased sampling of protein conformations. It is therefore a common practice to convert (i.e., reverse-mapping) an equilibrated CG Martini model to an AA models for simulating protein dynamics (2). For clarity, this hybrid CG-equilibrium-AA-production protocol will be referred to as “CG-to-AA” simulation.

The CG-to-AA strategy is especially useful for simulating large proteins such as mechanosensitive Piezo1 and Piezo2 channels (5, 6). A functional Piezo channel is formed by the assembly of three subunits, each encompassing ∼2500 amino acids. The homotrimeric Piezo channel structure displays a central pore and three highly curved transmembrane domains called arms (or blades). When embedded in a lipid bilayer, the Piezo arms curve the surrounding membrane into an inverted-dome shape, as evidenced from cryo-EM structures solved in detergent micelles and liposomes (7-11), and from AA (12) and CG-to-AA (13, 14) MD simulations performed in explicit solvent and membrane at body temperature. In particular, AA simulations have shown that the curvature mismatch between the curved (nonconducting) Piezo1 structure and the flat lipid bilayer drives the Piezo-membrane system into an equilibrium in which the Piezo1 arms become more flat and the membrane becomes more curved. This “tug-of-war” force balance between the protein and membrane took about 3 μs to reach equilibrium using AA simulation (12), but less than 300 ns using CG Martini simulation, thanks to the faster diffusion and smoothened free energy landscape of the CG model (13). Hence, CG-to-AA is computationally more efficient than AA alone for preparing large membrane proteins systems with complex topology.

Both Piezo1 arm flattening and pore dilation were observed using CG-to-AA simulations, either by imposing a protein-membrane curvature mismatch (using periodic boundary conditions) (13), or by lipid bilayer stretching (15). The open state obtained using curvature mismatch was further validated based on experimental conductance, selectivity, and mutant phenotypes (13). Remarkably, two important features predicted by the curvature mismatch simulation (clockwise rotation of the extracellular cap domain and dilation of the hydrophobic constriction site upon arm flattening) were perfectly recapitulated in a recent cryo-EM open state structure, also obtained using curvature mismatch (8).

A common observation from CG-to-AA simulations was the presence of several whole lipids (i.e., headgroups and tails) inside the nonconducting Piezo1 pore at the CG equilibrium stage. After reverse-mapping to AA model, those lipids either remained in the pore during three replicas of 2 μs AA simulations (13), or diffused away under large membrane tension (15). Lipid-mediated pore occlusion has emerged as a novel gating paradigm for mechanosensitive channels, such as the *E. coli* small-conductance mechanosensitive channel MscS (16) and the human TWIK-related arachidonic acid-activated potassium (TRAAK) channel (17). Hence, the mechanism by which these lipids enter the channel pore during MD simulations deserves close scrutiny, as these lipids may radically change our understanding of channel gating.

We began this study with a dilemma: the lipid density in the Piezo1 pore was found to be higher in the CG-to-AA system (using Martini v2.2 CG force field and CHARMM36 AA force field) relative to the AA system using the same AA force field. Yet, no cryo-EM electron density corresponding to lipid headgroups can be seen in the central pore (7-11). Furthermore, our AA Piezo1 simulation did not show permanent residence of pore lipids during 8 μs simulation (12). To solve this dilemma, we tested four equilibrium protocols using a Piezo1 nonconducting pore model and compared lipid headgroup, lipid tail, and water density among CG, CG-to-AA, and AA simulations. We found that the final outcome of AA simulations using the CG-to-AA strategy depends on the pore solvation algorithm and the type of lipid restraint used during CG equilibrium. Furthermore, we show that some pore lipids observed at the end of CG-to-AA trajectories exhibit a positive binding free energy, suggesting that these lipids were kinetically trapped due to the slow diffusion and shorter trajectory of the AA simulation. We found that using a whole-lipid restraint during the initial CG equilibrium helps alleviate such artifacts in CG-to-AA simulations.

## METHODS

### CG simulation setup and protocols

Atomistic model of nonconducting piezo1 pore (residue 1976-2546) was truncated from our previously Piezo1 model built from cryo-EM structure (PDB 6b3r) (13). The CG simulation was executed in GROMACS version 2016.4 simulation package with the standard Martini v2.2 force field parameter settings(4, 18). The Piezo1 pore CG models were embedded in a POPC bilayer using INSANE (INSert membrane) CG building tool (19). The overall workflow of the simulations included energy minimization, isothermal-isochoric (NVT) and isothermal-isobaric (NPT) equilibrium runs, and NPT production runs (20, 21). General protocols in each stage are provided in **Supporting Methods**.

During all CG simulations, positional restraints were applied to the protein backbone with a force constant of 10000 kJ/mol/nm^2^ to maintain the nonconducting Piezo1 pore conformation. In addition, two types of lipid restraints were tested: one, labelled as ‘headgroup-only restraint’, is only applying positional restraints on phosphate headgroup beads of POPC lipids, and another, labelled as ‘whole lipid restraint’, is restraining the whole POPC lipids. For each restraint type, four replicas of ‘slow-release’ and three replicas of ‘fast-release’ equilibrium protocols were carried out (details in **Table S1a and S1b**).

### AA simulation setup and protocols

CHARMM36 force field was used for all AA simulations regardless of the simulation engines. Piezo1 pore AA system was solvated with CHARMM TIP3P water (22) and 150 mM KCl using the CHARMM36 force field (23). GROMACS commend “gmx solvate” was used to add water molecules. Water molecules were deleted from the box if the distance between any atom of the solute molecule(s) and any atom of the solvent molecule is less than the sum of the scaled van der Waals radii of both atoms which is smaller than a standard water bead of 5 Å (24, 25). To compare with CG MD simulations, this AA system was first minimized using 5000 steepest descent cycles in GROMACS 2016.4 package (18), and then underwent six stages of thermal equilibrium phase at 310.15 K (details in **Supporting Methods** and **Table S2a**).

For AA simulations, two replicas of 1μs production were conducted using PEMED CUDA module of Amber18 packages (26) with positional restraint of 100 kcal/mol/Å^2^ on protein backbone (details in **Table S2b**). Other parameters are the same with GROMACS setting except the temperature control was done using Langevin thermostat with a gamma parameter (friction coefficient) of 1.0 ps^-1^, and pressure coupling was using a semi-isotropic Monte-Carlo barostat with a target pressure of the 1.0 bar. The SHAKE algorithm (27) was used to constrain bonds involving hydrogen. For CG-to-AA model, two replicas of 200 ns production run were performed in Amber18 packages.

### Absolute binding free energy calculation

The absolute binding free energy of lipids in channel pore corresponds to the thermodynamic reversible work to move the lipids from the bilayer to the binding site. Same as ligand-protein binding, this thermodynamic quantity can be calculated through potential of mean force (PMF) approach in which the lipid is physically pulled away from the binding site or alchemical approach in which the nonbonded interactions are slowly decoupled (28). In this work, we chose alchemical approach because it is practical to treat the cluster of three POPC lipids in the pore as a single ligand regardless of their order and pathway of entering the pore. FEP/ *λ* replica-exchange molecular dynamics (FEP/ *λ* -REMD) in NAMD2.14 was used (29, 30). Unlike conventional small-molecule ligands, lipids inside a channel pore can display larger mobility and conformational flexibility. To make sure that our FEP/*λ*-REMD samples the correct ensemble, 100ns unbiased trajectories were used to compute the distribution of (1) the distance R between center of mass of three lipids and the pore, and (2) lipid conformational RMSD after rigid-body alignment of the pore to the initial frame (called DBC restraint in NAMD Colvars (31). The upper-limit of the ensemble distribution of R and RMSD from unbiased simulations were used to setup the upper-boundary of the flat-bottom harmonic restraints for the lipid RMSD and for distance (force constant of 100 kcal/mol/Å^2^). **Figure S1** shows the fully bound lipids do not reach the upper boundary of the RMSD flat-bottom harmonic restraint during unbiased simulation and *λ* = 1 perturbation stage. Thus, the RMSD restraint only took effect at uncoupled state and intermediate states.

In addition, to ensure that FEP/*λ*-REMD sampled the two end states properly, lipids outside the pore were prevented from entering the pore during FEP so that water can enter the pore during the decoupling stage. This was achieved by a RMSD restraint on the center-of-mass of a lipid (selected atom names are N C2 C218 C316 in Charmm36 force field) between current frame and initial frame. **Figure S2** shows the higher number of water molecules and absence of lipid headgroup in the pore at fully uncoupled state (*λ* = 0) agree well with the AA unbiased simulation. Likewise, the lower number of water and three lipid headgroups in the pore are also consistent between fully couple FEP state (*λ* = 1) and the CG-to-AA simulation with headgroup-only restraint during the CG equilibrium stage. Therefore, the initial and final states of FEP capture the correct conformational ensemble of the true end states (bound vs. unbound) of unbiased simulations. The restraint details are further described in **Supporting Methods**. All NAMD input files are provided at https://github.com/reneejiang/pore-lipids.

A total number of 128 replicas were used for the binding site and 64 replicas for two bulk bilayers of size 40 and 60 lipids per leaflet. The “softcore” potentials were used to avoid end-point catastrophe (32, 33). Each replica in the FEP/ *λ*-REMD simulation represents a state along the coupling parameter, and periodic swap is attempted between neighboring replicas every 100 steps (0.2ps). The accuracy of FEP/ *λ*-REMD depends on the overlaps between two potential energy distributions, which can be reflected by the acceptance ratio between replicas. The acceptance ratios between each adjacent pair are between 40%-80% for 128 *λ* Piezo system and 30%-70% for 64 *λ* bilayer-only system (**Figure S3**). Convergence was monitored by the time dependence of each predicted free energy term. This sampling time dependence provides an asymptotically unbiased estimator for each Δ*G*. We considered the FEP/ *λ*-REMD simulation is converged when the running average of 1 ns free energy value fluctuates within 0.5kcal/mol (**Figure S4)**. The free energy contribution of each term is listed in **Table S3**. The uncertainties were computed from pymbar (34, 35).

### Analysis of atoms/beads at Piezo1 pore regions

The counting method for classifying atoms/beads inside pore regions was based on the MATLAB function “inpolyhedron” (36) which can efficiently classify whether a point inside of a 3D triangulated surface. By adding surfaces to build up a closed volume for upper pore region, hydrophobic constriction site and lower pore region along the z-axis, the time series of the number of AA atoms or CG beads inside each region were classified and plotted. An example plot of the classification method is shown in **Figure S5**. The MATLAB codes are provided at https://github.com/reneejiang/pore-lipids.

## RESULTS

To investigate lipid density inside the central pore of Piezo1 channel, we constructed a pore model in the resting state (PDB 6b3r), which includes the cap domain, repeat A, anchor, pore, and CTD domains (residue V1976 to R2546, **Figure 1a**) (7). Previous MD simulations have shown that widening of a constriction site, consisting of three V2476 residues, is required for the Piezo1 pore to conduct ions with conductance and cation selectivity consistent with experimental measurements (13). Hence, to quantify the number of lipid headgroups, tails, and water molecules inside the pore during simulations, we define the “upper pore region”, “hydrophobic constriction site” and “lower pore region” by the positions of P2455 on the linkers between the cap and TM38, L2469, V2476 and F2485 on the inner pore helices (TM38) (**Figure 1b**). In all simulations, the protein backbone was constrained to the original cryo-EM conformation to ensure a fair comparison of lipid/water densities among simulations. In total, fourteen CG-MD simulations using Martini v2.2 force field and four AA-MD simulations with CHARMM36 force field were conducted to compare the lipid and water density in the central pore (**Figure 2** and **Table S1**).

**Figure 1.**
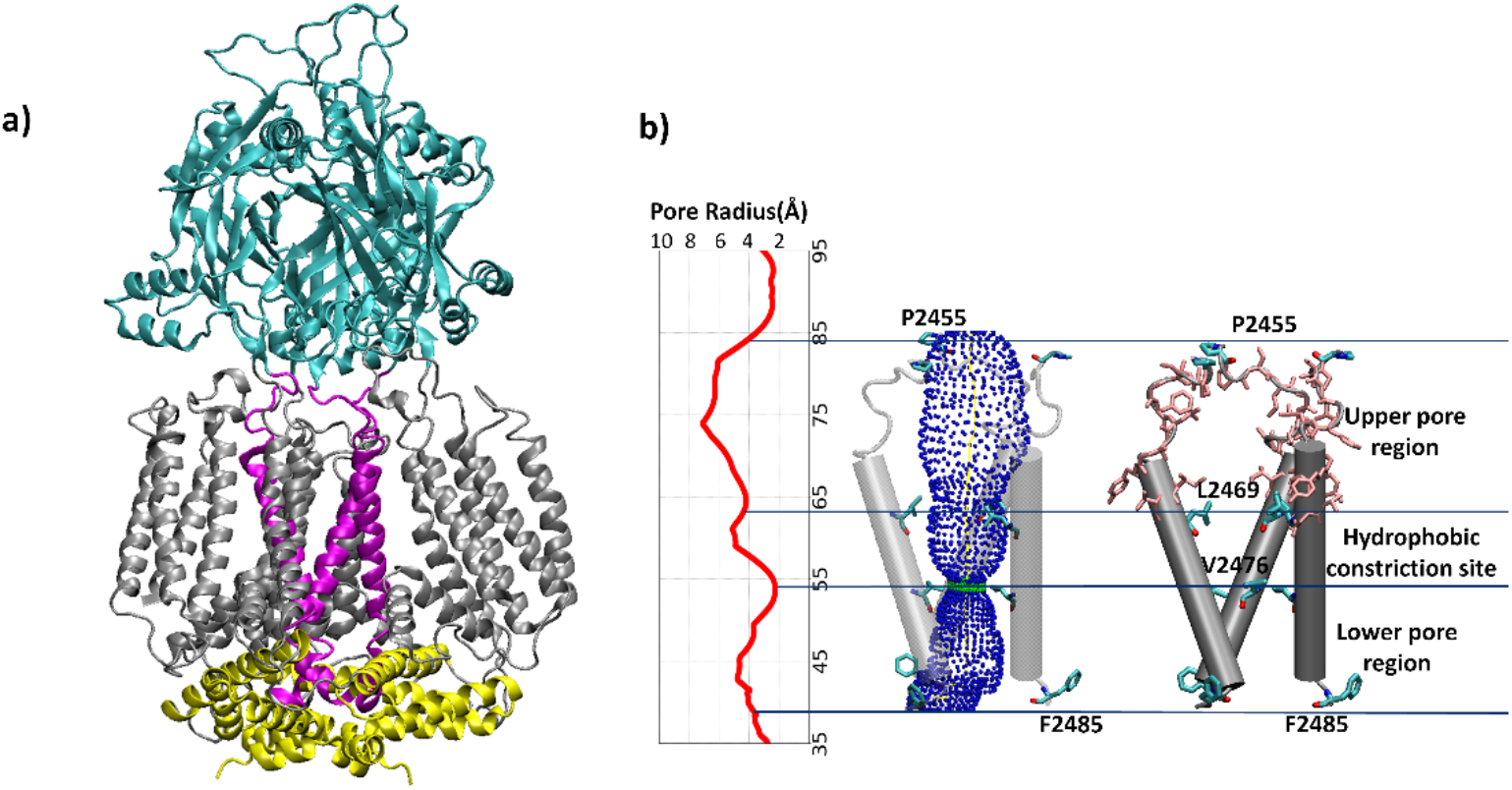
Piezo1 pore model and the definition of the upper and lower pore regions. **a)** The Piezo1 pore model (residue V1976 to R2546) is mainly composed of the cap colored in cyan, pore in purple, CTD region in yellow, and repeat A in silver. **b)** Pore radius profile for the cryo-EM solved Piezo1 pore and classified regions defined to count the number of water and lipid molecules inside. The residues P2455, L2469, V2476 and F2485 are in licorice mode. Pore shown in 3D is calculated using HOLE program (37) and plotted in VMD (38). The pink residues in licorice mode are the hydrophobic residues along the Piezo1 upper pore region.

**Figure 2.**
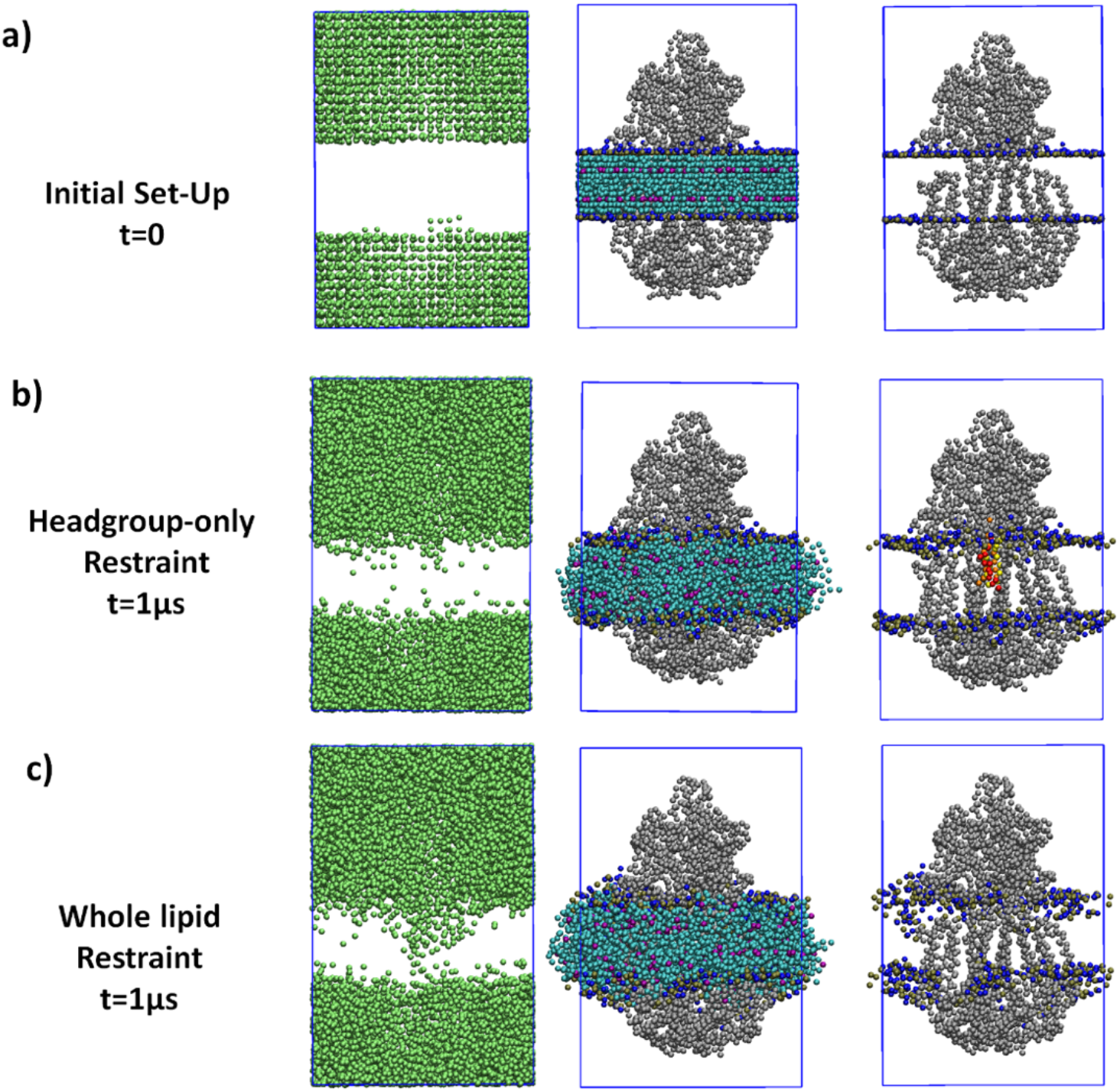
Illustration of water and lipid beads in Piezo1 pore domain from CG simulations. **a)** CG initial system setup by components and a close look of pore region. **b)** System with lipid headgroup-only restraint during equilibrium stage and fully released bilayer for 1μs production run. **c)** System with whole-lipid restraint during equilibrium stage and fully released bilayer for the 1μs production run. Color scheme: water beads in green, protein backbone in silver, lipid phosphate group in brown, lipid nitrogen bead in blue, lipid tail beads in cyan and purple. The 3 lipids trapped during reverse-mapped AA simulations are colored in red, yellow, and orange. All the beads are shown in VDW mode in VMD.

### Fast vs. slow release of lipid headgroup restraint in CG model

During the Martini CG bilayer system setup using insane.py, solvent molecules were generated using a 3D grid where the grid cells occupied by membrane and/or proteins were flagged unavailable, and the remaining cells were filled with solvents (19). In the default setting, water molecules are added above and below the bilayer, thus left the pore region empty (**Figure 2a**). This solvation step is followed by equilibrium steps, in which water molecules first undergo equilibrium when protein beads and lipid headgroup beads are restrained. To investigate the effect of this equilibrium protocol on the lipid density inside the pore, we tested a ‘fast-release’ and a ‘slow-release’ of lipid headgroup restraints.

In the ‘fast-release’ protocol, the headgroup force constant was reduced from 5000 to 10kJ/mol/nm^2^ during 10 ns, and from 10 to 0 kJ/mol/nm^2^ during 5 ns (**Table S1a**). During three replicas of ‘fast-release’ equilibrium runs, 2-3 lipids headgroups rapidly moved into the center and remained during 1 μs CG production runs (**Figure 2b, 3a**). There are also 2 to 3 lipid tails (one CG POPC lipid model contains 8 tail beads) remained in the upper pore region (**Figure 3c**). Consequently, water molecules are largely excluded from this pore region (**Figure 3b**). In contrast, four replicas of ‘slow-release’ protocol with the headgroup force constant reduced from 5000 to 1000kJ/mol/nm^2^ during 5 ns (**Table S1b**) showed fewer lipids in the center region (1-2 headgroups vs. 2-3 in the ‘fast-release’ runs), and more water molecules in the pore (**Figure 3d-f**). Overall, our data suggest that the vicinity of lipids near the hollow pore could allow unrestrained lipid tails to enter the pore before water molecules through spaces formed between neighboring pore helices. Once the lipid headgroup restraint is removed, those lipid tails quickly pulled their headgroups inside the pore, altogether preventing hydration of the pore. Slowly releasing the headgroup restraint did not completely eliminate this artifact.

**Figure 3.**
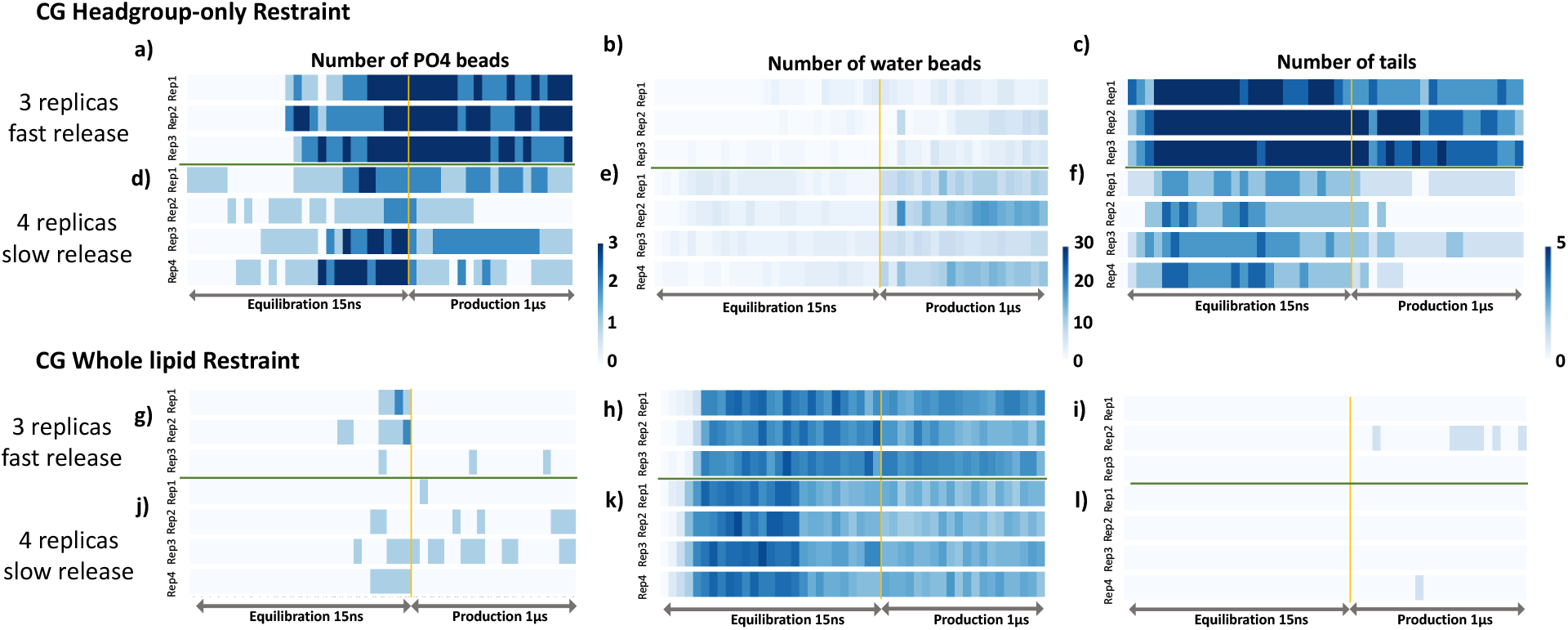
Number of CG lipid headgroup PO4 beads, water beads, and tails in Piezo1 upper pore region during equilibrium and production runs. In Martini, four water molecules are mapped into one bead, one POPC lipid has 1 headgroup bead PO4 and 8 tail beads. ‘Headgroup-only restraint’ and ‘Whole lipid restraint’ both include the ‘fast-release’ (**Table S1a**, replica 1-3) and ‘slow-release’ equilibrium protocol (**Table S1b**, replica 1-4) for the bilayer during equilibrium stage (see **Table S1** for details of equilibrium protocols). The production stage ran for 1μs for both CG models with 10000 kJ/mol/nm2 position restraints on protein backbone. Yellow line separates the equilibrium and production stages. Green line separates the ‘fast-release’ and ‘slow-release’ equilibrium protocols.

### Fast vs. slow release of whole-lipid restraint in Piezo1 CG model

The seven aforementioned CG simulations (**Figure 3a-f**) show that lipids prevent pore hydration by infiltrating the hollow pore. We thus asked if restraining the whole lipid molecules instead of the headgroups during CG equilibrium would promote pore hydration. Using a whole lipid restraint, water beads were able to enter the pore and the pore remains hydrated in both fast and slow-release protocols (**Figure 3g-l**). During the production run, both lipid headgroups and tails could enter the pore sporadically but did not reside in the pore to the same extent as in the CG headgroup-only restraint simulations. In conclusion, the whole lipid restraint setting allowed ample time for water molecules to diffuse into the upper pore region during equilibrium stage, while in the headgroup-only restraint, lipids were able to pre-occupy the pore, reducing pore accessibility to water molecules throughout the subsequent CG simulations. This effect is particularly noticeable when the headgroup-only restraint is released quickly. Implementing the whole-lipid restraint reduces the effect of the speed of restraint release on the production run and led to a hydrated pore that agrees better with the system prepared directly from AA force field below.

### Pore hydration during AA equilibrium

The default GROMACS solvation algorithm (25) places water molecules in the empty space of bilayer and protein region (**Figure 4a**), in contrast to the default insane.py setting in CG model (19), which placed no water beads in bilayer and protein regions (**Figure 2a**). Using lipid headgroup-only restraint at AA equilibrium stage (detailed in **Table S2a**), we found no lipids accumulated at the upper pore region during two replicas of 1 μs production run (**Figure 4b and 5a**). The water density plot shows that both upper and lower pore regions are hydrated, with a near-zero water density around the hydrophobic constriction site, as expected. This data is consistent with our previous 8 μs AA simulation of Piezo1 nonconducting state prepared using CHARMM-GUI protocol, which also adds water inside the channel pore during the solvation step (12).

**Figure 4.**
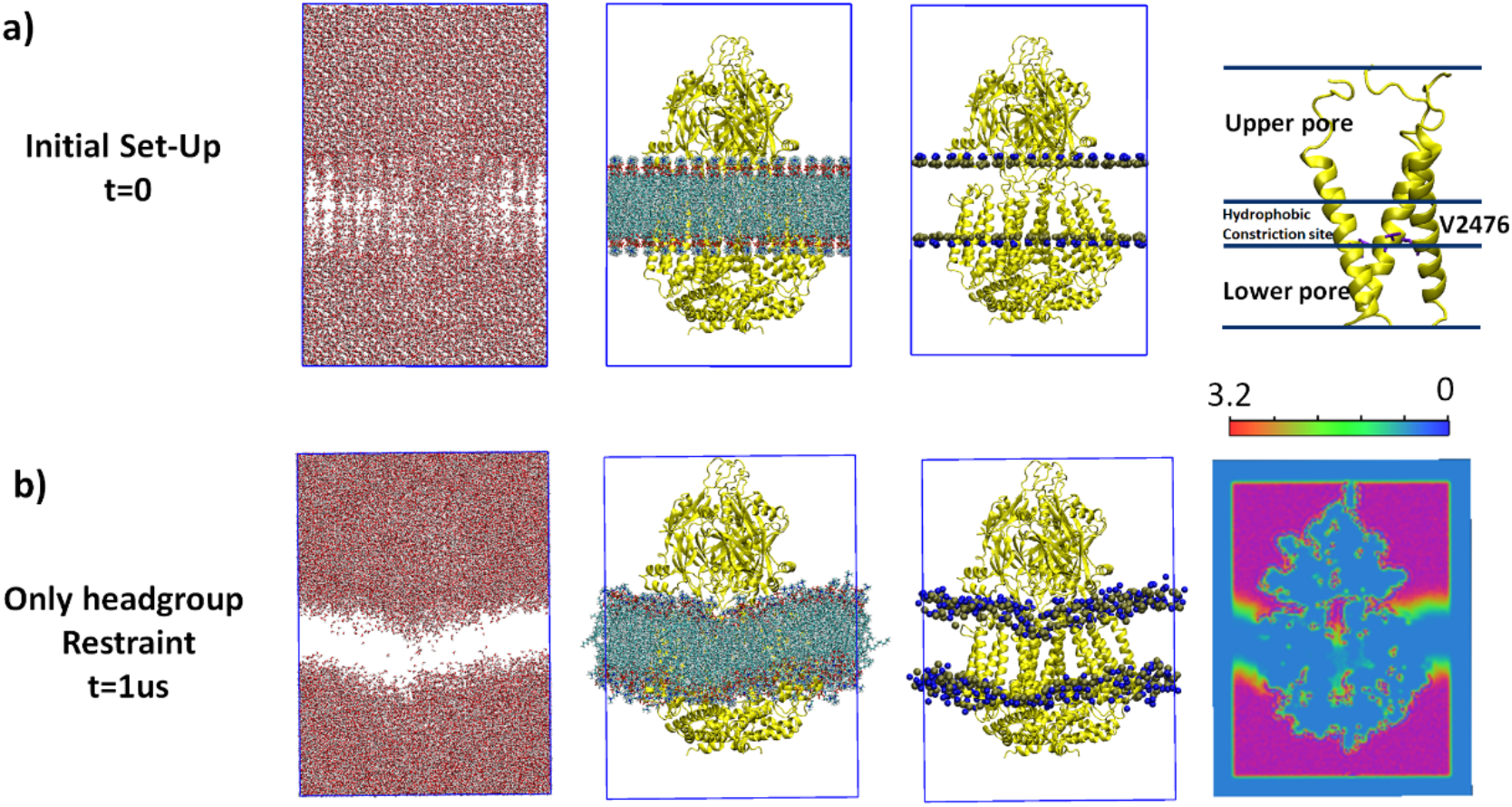
AA simulation of Piezo1 pore domain. **a)** AA initial system setup by components, and a close look of pore region divided into upper pore region, hydrophobic constriction site and lower pore region. **b)** Following the standard CHARMM-GUI protocol (using headgroup-only restraint) during the equilibrium stage and fully release the restraint but protein backbone for the 1μs production stage. Color scheme: oxygen atom in red, hydrogen atom in white, protein in yellow, V2476 residue in violet, phosphorus in brown, nitrogen in blue, hydrophobic tails in cyan. Water and lipids are shown in VMD licorice mode, the third column showing phosphorus and nitrogen atoms in VDW mode, protein is shown in new cartoon mode. The water density map is computed from the last 500ns for AA system and visualized with slice offset=0.46 along *y*-axis in VMD.

**Figure 5.**
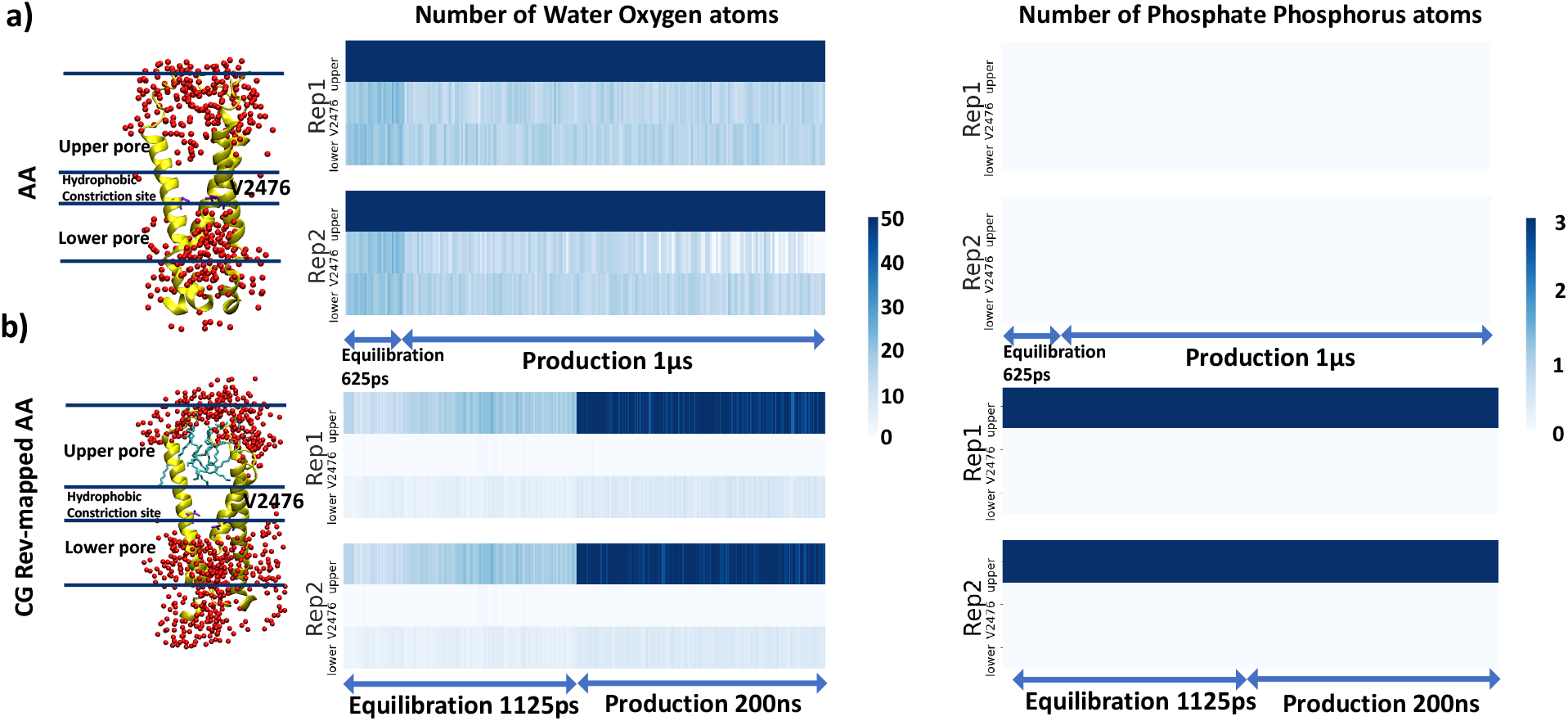
Number of water oxygen and lipid headgroup phosphorus atoms in upper pore region of the AA model and CG Rev-mapped AA model. Both two AA systems used the standard CHARMM-GUI equilibrium protocol with only lipid headgroup restraint for the bilayer during equilibrium stage, and fully released bilayer during the production stage. The production stage ran for 1μs for AA model and 200ns for CG Rev-mapped AA model with 10000 kJ/mol/nm2 position restraints on protein backbone. Each restraint system ran for two replicas with the same protocol (see **Table S2a and S2b** for details of the standard equilibrium protocol). Color scheme: oxygen atom in red, V2476 residue in violet, protein in yellow, lipids in cyan, all other components are not shown here.

### Comparing AA model with the CG Rev-mapped AA model

We next asked whether reverse-mapping the CG model with lipids in the pore to the same AA force field would converge to the similar hydration pattern as seen in the AA model. Two replicas of fast-release CG headgroup-only restraint system **(Figure 3a)** were reverse-mapped back to AA system (CG-to-AA model), and the AA production run was extended for 200 ns to compare the number of water molecules and lipid headgroups in the upper pore region with that observed in the original AA model. **Figure 5b** shows that the CG Rev-mapped AA model, which initially contained 20 water molecules during the equilibrium stage, contains 46 water molecules during the production stage, with all 3 initial lipids remaining inside the upper pore region during the 200 ns production run. In contrast, in both replicas of the original AA model, the upper pore remains hydrated by more than 50 water molecules and no lipid headgroup were present in the pore (**Figure 5a**).

### Comparing lipid distribution from AA simulation with cryo-EM lipid density

While taking a close look at the lipids around Piezo1 pore in our AA simulation (**Figure 5a**), we find lipids lining the hollow space formed between neighboring outer helices and inner helices (pink and yellow helices in **Figure 6a**). We define these lipids as ‘wall lipids’. During 1 μs AA production run, between 3 to 11 lipid tail carbon atoms are seen entering the pore sporadically through spaces between inner pore helices (**Figure 6a and S6**). However, no full lipid was seen occluding the pore. In contrast, in CG-to-AA simulations (**Figure 5b**), in addition to the wall lipids, 3 full lipids occupy the space below the Piezo1 cap domain and above the hydrophobic constriction site of the pore (**Figure 6b**). While lipid tails are seen to fluctuate in and out of the upper and lower pore region frequently, lipid headgroups are clearly trapped in the upper pore throughout the simulations. We define these lipids as ‘pore lipids’. A recent cryo-EM structure of non-conducting Piezo1 in small liposomes confirmed the ‘wall lipid’ density seen from the lower leaflet, with lipid tails extending into the lower pore and headgroups sitting above the lower fenestration (**Figure 6c**), but no lipid density was captured from the upper leaflet or along the central axis of the pore (8), consistent with our AA simulation results.

**Figure 6.**
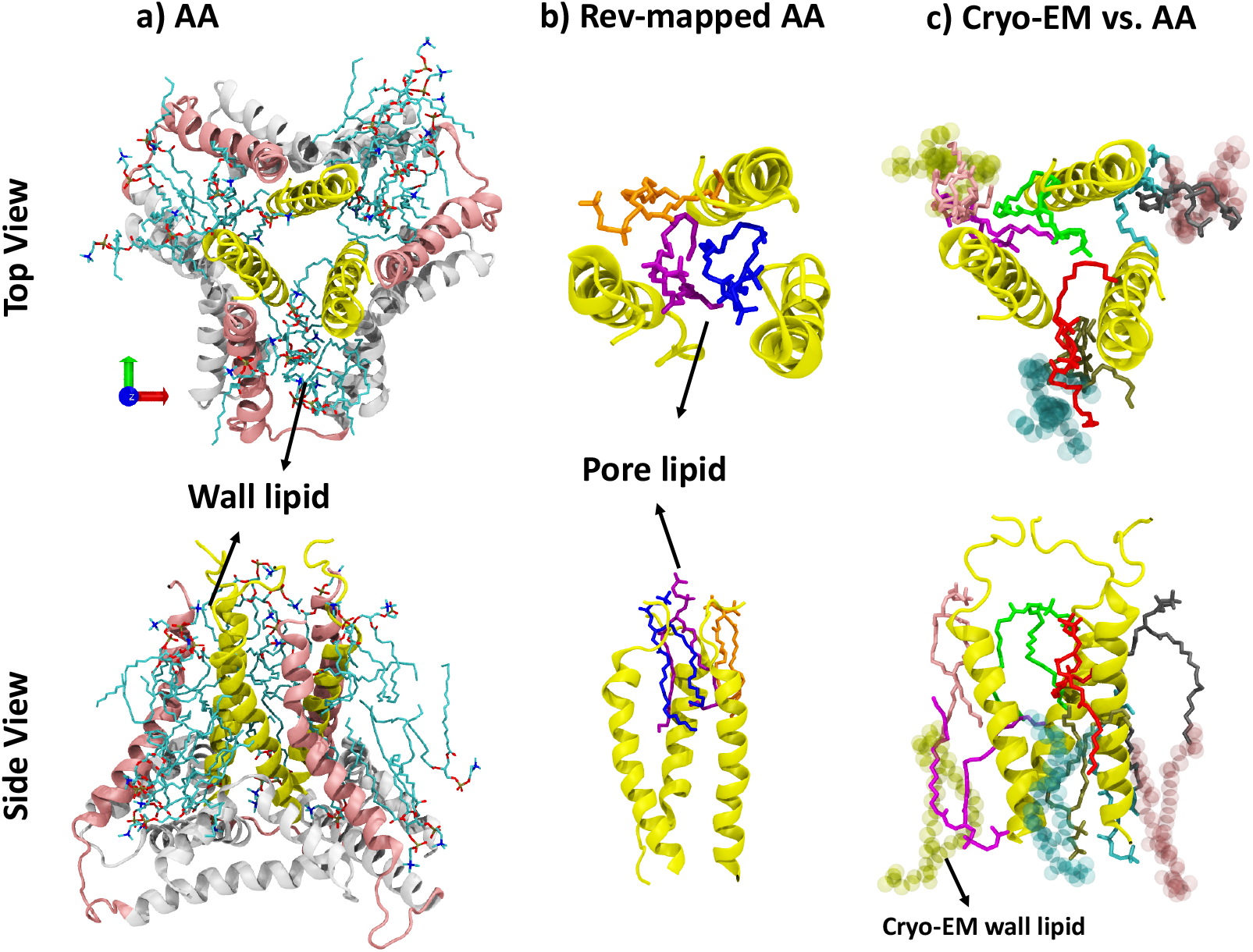
**Top view and side view of lipids around the Piezo1 pore** for AA, CG Rev-mapped AA and Cryo-EM Piezo1 (PDB ID: 7wlt) model. **a) AA**: yellow in new cartoon is inner helix, pink is outer helix, white is the anchor region, oxygen in red, phosphors in brown, nitrogen in blue, carbon in cyan. **b) Rev-mapped AA**: three trapped lipids are in blue, orange, and purple color. The CG Rev-mapped pore helix have a bit of twist due to the process of reverse-mapping from CG to AA model. **c) Cryo-EM Piezo1**: lipids from Cryo-EM structure are in VDM mode with transparent color of yellow, cyan, and pink. Lipids from AA model simulations are in licorice mode.

### Absolute binding free energy of lipids in Piezo1 pore

Are these POPC lipids inside nonconducting Piezo1 pore (**Figure 6b**) energetically stable? To answer this question, we sought to compute the absolute binding free energy of pore lipids, starting from the final configuration of the CG-to-AA in which three full lipids remain in the center of the upper pore region for 200 ns (**Figure 5b**). Here, absolute free energy is defined as the free energy difference between lipids in the pore and lipids in a homogenous POPC bilayer (**Figure 7a**). FEP/ *λ* replica-exchange molecular dynamics (FEP/ *λ* -REMD) in NAMD2.14 was used. In order to sample the high flexibility and mobility of POPC lipid’s long fatty acid aliphatic chains, we adopted the flat-bottom harmonic restraints on the perturbed lipids inside the pore and in bilayer (31) (**Figure 7b**). This restraint protocol allows unbiased sampling of lipids in the binding site when they are fully coupled (**Figure S1**), and is thus more suitable to highly mobile ligands or lipids than the Boresch-style restraints (i.e., six rigid-body restraints) (39) commonly applied to small drug molecule binding. The detailed methods and convergence analysis are provided in **Methods** section. Our results indeed reveal that a full lipid in the pore is highly unfavorable (+10.0 ± 0.8 kcal/mol), which suggests the lipids are kinetically trapped in the pore during AA simulation.

**Figure 7.**
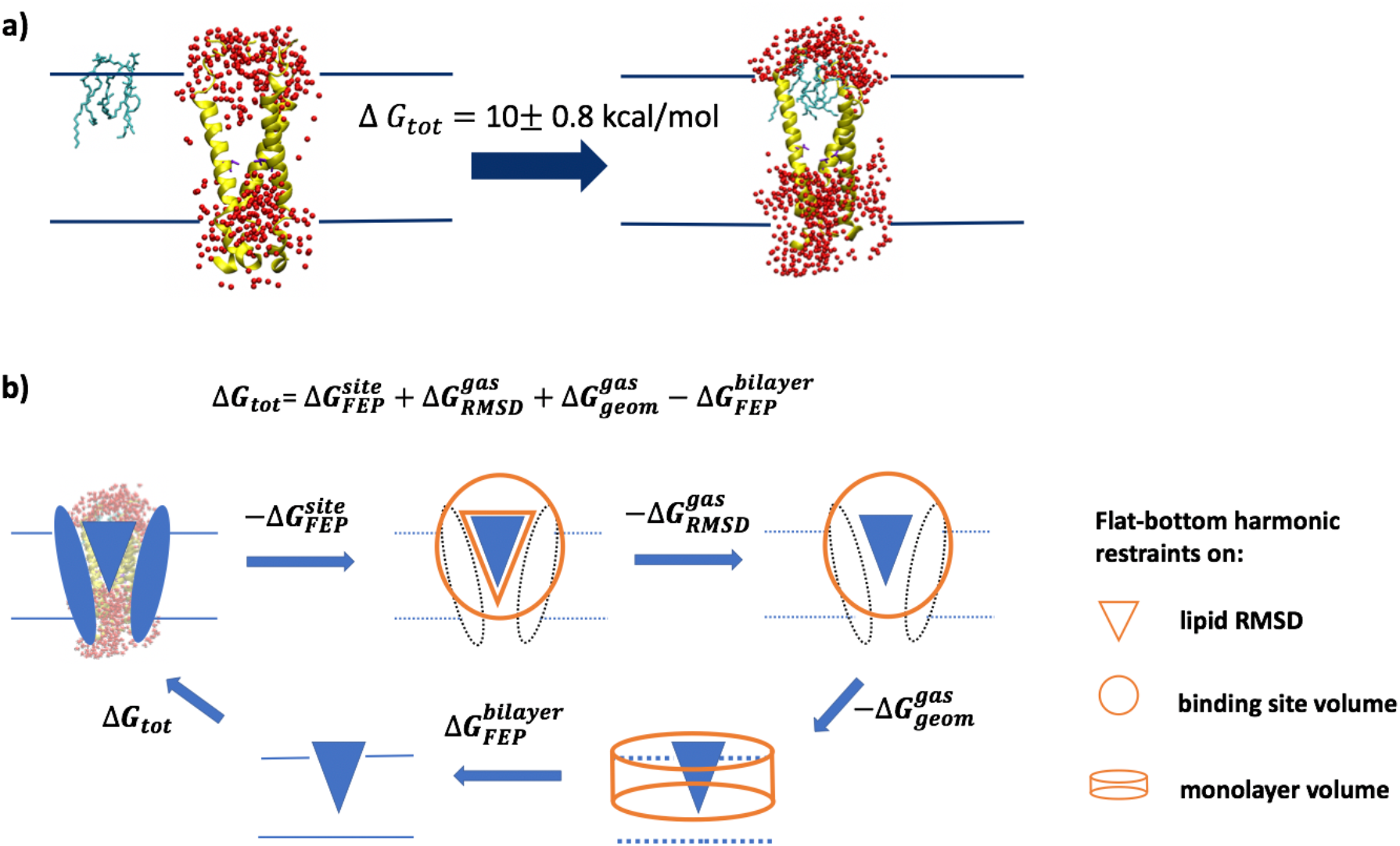
Binding free energy calculation of lipids in Piezo1 pore. **a)** The absolute binding free energy of lipids in Piezo1 pore corresponds to the reversible work to move the lipids from the bilayer to the pore. Color scheme: water oxygen atom in red, pore helix in yellow, and pore lipids in cyan. All other components are not shown here. **b)** Graphic presentation of the thermodynamic circle of FEP protocol (see **Supporting Methods** and **Table S3**).

## DISCUSSION

Protein-lipid interactions have drawn unprecedented attention in recent years, thanks to the increasing number of high-resolution membrane protein structures solved in a lipid environment (40). The identification of lipid-like electron densities in these structures, coupled with functional studies, constitutes a powerful experimental approach to investigate the contribution of lipids to channel gating (41). CG simulations using Martini force field has been successful in capturing lipid-protein interaction dynamics (42) and computing lipid binding free energy and kinetics (28, 43). An obvious advantage of CG Martini model is that the diffusion coefficient of CG lipid models are at least four times faster than in AA models (42). Thus, it often goes unquestioned that a CG-equilibrium-AA-production protocol is an efficient and reliable way to prepare membrane protein systems before investigating protein dynamics with an atomistic resolution. Here, we show that this hybrid CG-to-AA protocol can result in biased lipid density in a channel pore.

Discrepancy in pore lipid density have been reported by structural biologists when comparing different protein purification and reconstitution approaches (44). Here, the discrepancy in pore lipid density from MD simulations is partially due to the topology of the homotrimeric Piezo1. The three inner pore helices form an hourglass shape with a hydrophobic constriction site formed by three V2476, and three fenestrations above and below this constriction site (13) (**Figure 1**). In CG model preparation, the protein is first embedded in a bilayer and then water beads are added above and below the bilayer, but not in transmembrane protein crevices such as channel pores. In the equilibrium step, both the protein and lipid headgroups are restrained to allow water molecules to diffuse freely and to fill in empty space. In principle, this setup should allow full hydration of the pore if the pore walls are sealed against lipids. However, for channels whose pores are walled with transmembrane gaps, adjacent lipids near the gap may enter the pore before the pore gets fully hydrated. This is partially due to the faster diffusion of CG Martini lipid model and larger CG water beads. Once the CG system is reverse-mapped back to AA system, lipids are trapped in the pore because AA lipids diffuse much slower and free energy surface in AA model is less smooth than in CG model.

In conclusion, the geometry of the pore, the lack of initial pore hydration, and the mismatch between fast CG and slow AA model all play a role in this “trapped pore lipids” artifact. This artifact can be easily eliminated by restraining whole lipids during initial water diffusion. Indeed, this artifact is not seen in our tested AA model because the AA solvation algorithm adds water in the pore before the equilibrium step. By restraining whole lipids during CG equilibrium, we obtain similar results as those obtained using AA equilibrium, including our previous 8 μs-long simulation (12), that is, the absence of whole lipids in the pore. Our simulation is further confirmed by the lipid density reported in a recent Piezo1 cryo-EM curved conformation, in which the lower leaflet “wall lipids” enter the pore through the crevices between inner pore helices, but do not block the pore entirely (**Figure 6c**).

This work by no means excludes the contribution of lipids to Piezo1 function. In fact, it provides a more robust starting point for further investigation. For instance, the cryo-EM lipid density captured near the lower fenestration of Piezo1 pore (8) could play a role in channel gating. It is also possible that when the pore dilates during Piezo1 activation, the crevices between inner helices become large enough to allow whole lipids to enter and block the pore. While we only focused on the equilibrium protocol of CG simulations in this study, the comparison of binding free energy of lipids in the Piezo1 open pore in different models, such as CG, AA, or hybrid resolution models, needs further investigation. In addition, although we only studied POPC lipids here, this work should serve as a benchmark for future investigations of more complex Piezo-lipid interactions in heterogenous bilayers. Indeed, many lipids are known to modulate Piezo1 function such as PIP_2_(45), PS (46), margaric acid-enriched phospholipids (47, 48), and sterols (49-51). Understanding the role of lipids to the function of mechanosensitive ion channels remains a central question in the field of mechanobiology. Eliminating possible computational artifacts is an important step toward achieving this goal.

## CODE AVAILABILITY

Input files and coordinates of Piezo1 system for MD simulations and free energy calculations, as well as python2 scripts and MATLAB codes are publicly available at https://github.com/reneejiang/pore-lipids.

## ACKNOWLEDGMENTS

This work was supported by NIH Grants GM130834 (Y.L. and J.L.). Computational resources were provided via the Extreme Science and Engineering Discovery Environment (XSEDE) allocation TG-MCB160119, which is supported by NSF grant number ACI-154862. Anton2 computer time was provided by the Pittsburgh Supercomputing Center (PSC) through NIH Grant R01-GM116961. The Anton2 machine at PSC was generously made available by D.E.Shaw Research.

## AUTHOR CONTRIBUTIONS

W.J. performed all MD simulations and data analysis. Y.L designed and supervised the project. W.J., and Y.L and J.L wrote the paper together.

