## Supplemental material for "Importance of Molecular Dynamics Equilibrium Protocol on Protein-lipid Interactions near Channel Pore"

### Supporting Methods

#### CG simulation protocols

A cut-off of 1.2 nm was used for calculating both the electrostatic and van der Waals interaction terms; the potential-shift-Verlet algorithm was applied to take care of both interactions by smoothly shifting beyond the cutoff. Coulomb interactions were calculated using the reaction-field algorithm implemented in GROMACS. The neighbor list was updated every 20 steps using a neighbor list cutoff equal to 1.2 nm for short-range van der Waals. The temperature for each group (protein, membrane, ion, and water) was kept constant at 310.15 K using the velocity rescale coupling algorithm with 1 ps time constant. In the NPT equilibrium step, semi-isotropic pressure coupling was applied using the Parrinello-Rahman algorithm, with a pressure of 1 bar independently in the cross-section of the membrane and perpendicular to the membrane with the compressibility of $3.0 \times 10^{-4}$ bar$^{-1}$. The relaxation time constant for pressure coupling was set to 10.0 ps. Three-dimensional periodic boundary conditions were used. The CG system was first minimized using 5000 steepest descent cycles. Following minimization, in the first equilibrium stage, a 1250 ps NVT solvent heating simulation ran to attain a target temperature of 310.15 K with a timestep of 10 fs. A time step of 5 fs was used for the following NPT simulations due to high positional restraint on protein backbone.

**Table S1a**. **'Fast-release' protocol used for equilibrating the CG systems in GROMACS package.**

| Step[a] | Model Type | 6.1 | 6.2 | 6.3 | 6.4 | 6.5 | 6.6 | 7 |
|---|---|---|---|---|---|---|---|---|
| Protein Positional restraints[b] (kJ/mol/nm$^2$) | CG | 10000 | 10000 | 10000 | 10000 | 10000 | 10000 | 10000 |
| Lipid headgroup restraints[c] (kJ/mol/nm$^2$) | CG | 5000 | 5000 | 1000 | 1000 | 10 | 0 | 0 |
| whole lipid restraints[d] (kJ/mol/nm$^2$) | CG | 5000 | 5000 | 1000 | 1000 | 10 | 0 | 0 |
| Length of simulations(ns)[e] | CG | 1.25 | 1.25 | 1.25 | 1.25 | 5.0 | 5.0 | 1000 |

[a]Equilibrium steps include step6.1 to step6.6 and step7 is the production run

[b]Positional restrains on all the backbone beads in the protein

[c]Positional restraints only on the phosphate group PO4 bead of lipid POPC
[d]Positional restraints on each bead of the whole lipid POPC during the simulation
[e]Replica 1 to 3, referred as 'fast-release'

**Table S1b**. **'Slow-release' protocol used for equilibrating the CG systems in GROMACS package.**

| Step[a] | Model Type | 6.1 | 6.2 | 6.3 | 6.4 | 6.5 | 6.6 | 7 |
|---|---|---|---|---|---|---|---|---|
| Protein Positional restraints[b] (kJ/mol/nm$^2$) | CG | 10000 | 10000 | 10000 | 10000 | 10000 | 10000 | 10000 |
| Lipid headgroup restraints[c] (kJ/mol/nm$^2$) | CG | 5000 | 5000 | 3000 | 3000 | 1000 | 0 | 0 |
| whole lipid restraints[d] (kJ/mol/nm$^2$) | CG | 5000 | 5000 | 3000 | 3000 | 1000 | 0 | 0 |
| Length of simulations(ns)[e] | CG | 1.25 | 1.25 | 1.25 | 1.25 | 5.0 | 5.0 | 1000 |

[a]Equilibrium steps include step 6.1 to step 6.6 and step 7 is the production run
[b]Positional restrains on all the backbone beads in the protein
[c]Positional restraints only on the phosphate group PO4 bead of lipid POPC
[d]Positional restraints on each bead of the whole lipid POPC during the simulation
[e]Replica 1 to 4, referred as 'slow-release'

**AA simulation protocols**

All the stages and in both AA model and CG-to-AA model, a non-bonded cut-off 1.2 nm was employed for both the electrostatic and van der Waals interaction terms. The potential-shift-Verlet algorithm was applied to take care of both interactions by smoothly shifting beyond the cutoff. Coulomb interactions were calculated using PME algorithm implemented in GROMACS. Positional restraints were applied to backbone of protein residues with a force constant of 10000 kJ/mol/nm$^2$ over all the stages (details in **Table S2**. For all simulation stages, the neighbor list was updated every 20 steps using a neighbor list cutoff equal to 1.2 nm for short-range van der Waals and temperature control was accomplished using velocity rescale coupling algorithm with 1 ps time constant. The LINCS algorithm was used to constrain bonds involving hydrogen(1). For all AA simulations, the neighbor list was updated every 20 steps using a neighbor list cutoff equal to 1.2 nm for short-range van der Waals and temperature control was accomplished using velocity rescale coupling algorithm with 1 ps time constant. The LINCS algorithm was used to constrain bonds involving hydrogen.

In the NPT equilibrium step, pressure regulation was accomplished by utilization of a semi-isotropic Parrinello-Rahman barostat with a pressure of 1 bar independently in the cross-section of the membrane and perpendicular to the membrane with the compressibility of $4.5 \times 10^{-5}$ bar$^{-1}$. The relaxation time constant for pressure coupling was set to 5.0 ps. Following minimization, in the first equilibrium stage, a 125ps NVT solvent heating simulation ran to attain a target temperature of 310.15K. Phosphate atoms of membrane POPC residues were restrained with a force constant of 5000 kJ/mol/ nm$^2$. Dihedral restraints were also employed to ensure retention of backbone phi-psi angles during minimization with 2000 kJ/mol/radian$^2$ force constant. The second equilibrium stage followed an identical setup of 125ps with the sole exception that the force constant of the dihedral phi-psi restraint was also reduced to 1000 kJ/mol/radian$^2$. Starting with the third equilibrium stage, an NPT ensemble was employed. For the third stage of 125ps, force constants for phosphate atoms positional restraints were reduced to 3000 kJ/mol/nm$^2$. The dihedral phi-psi restraints were also dropped to 500 kJ/mol/radian$^2$. In the fourth stage, positional restraint force constants for phosphate atoms were maintained at 3000 kJ/mol/nm$^2$ and phi-psi backbone dihedral restraint force constants were reduced to 200 kJ/mol/radian$^2$. From the fourth to sixth stages, AA model ran for 125ps each and reverse-mapped AA model for 250ps, respectively. In the fifth stage, positional restraint force constants for phosphate atoms were reduced to 1000 kJ/mol/nm$^2$ and phi-psi backbone dihedral restraint force constants were reduced to 100 kJ/mol/radian$^2$. In the sixth equilibrium stage, membrane positional restraints and phi-psi backbone dihedral restraints were removed entirely. Simulation timestep was set as 1.0 fs due to high positional restraint on protein backbone.

**Table S2a**. **Multi-step procedure used for equilibrating the AA model in GROMACS package.**

| Step[a] | Model Type | 6.1 | 6.2 | 6.3 | 6.4 | 6.5 | 6.6 |
|---|---|---|---|---|---|---|---|
| Protein positional restraints[b] (kJ/mol/nm$^2$) | AA/Rev-AA | 10000 | 10000 | 10000 | 10000 | 10000 | 10000 |
| Dihedral phi-psi restraints[c] (kJ/mol/radian$^2$) | AA/Rev-AA | 2000 | 1000 | 500 | 200 | 100 | 100 |
| Lipid headgroup restraints[d] (kJ/mol/nm$^2$) | AA/Rev-AA | 5000 | 5000 | 3000 | 3000 | 1000 | 0 |
| Length of simulations(ns) | AA | 0.125 | 0.125 | 0.125 | 0.125 | 0.125 | 0.125 |
| | Rev-AA | 0.125 | 0.125 | 0.125 | 0.250 | 0.250 | 0.250 |

[a]Equilibrium steps include step 6.1 to step 6.6
[b]Positional restrains on all the backbone atoms in the protein
[c]Dihedral phi-psi restraints only on the protein atoms
[d]Positional restraints only on the phosphate group atom P in AA simulation

**Table S2b**. **Production run used for the AA model in Amber18 package.**

| Production Run | Model Type | step7 |
|---|---|---|
| Protein positional restraints[a] (kcal/mol/Å$^2$) | AA/Rev-AA | 100 |
| Lipid headgroup restraints[b] (kcal/mol/Å$^2$) | AA/Rev-AA | 0 |
| Length of simulations(ns) | AA | 1000 |
| | Rev-AA | 200 |

[a]Positional restrains on all the backbone atoms in the protein
[b]Positional restraints only on the phosphate group atom P in AA simulation

**Free Energy Perturbation Protocol for Lipid-Protein Binding**

Shown in **Figure 7**, the absolute binding free energy can be obtained using $\Delta G_{tot}$ = $\Delta G_{FEP}^{site} + \Delta G_{RMSD}^{gas} + \Delta G_{geom}^{gas} - \Delta G_{FEP}^{bilayer}$, where $\Delta G_{FEP}^{site}$ denotes the contribution of decoupling bound lipids ($\lambda = 1 \to 0$). A total of 128 $\lambda$ intermediates was used to ensure >40% acceptance ratio between neighboring states. $\Delta G_{RMSD}^{gas}$ denotes the contribution of flat-bottom harmonic RMSD restraint on conformational ensemble of lipids when lipids are fully uncoupled (i.e. in gas phase, $\lambda = 0$). $\Delta G_{RMSD}^{gas}$ was calculated using NAMD Colvars module(2) with 20 $\lambda$ values from 1.0 to 0.0 ( $\lambda = 1, 0.9999, 0.999, 0.99, 0.9, 0.8, 0.7, 0.6, 0.5, 0.4, 0.35, 0.325, 0.3, 0.25, 0.225, 0.2, 0.175, 0.15, 0.125, 0$ ). Each $\lambda$ was run for 200ps and errors were calculated from three replicas. $\Delta G_{geom}^{gas}$ denotes the entropic contribution, thus the volume ratio of fully uncoupled lipids in the binding site ($\frac{3}{4}\pi R^3$) versus the sampled volume of uncoupled POPC in the bulk bilayer. $R$ is the upper boundary of flat-bottom distance restraint between lipids and Piezo1 pore. In a bulk bilayer with surface area $A$, a flat-bottom restraint was introduced on the $z$-axis distance of the POPC lipid headgroup COM from the upper leaflet midplane $Z_R$ so that the uncoupled POPC remains in the upper leaflet. In addition, an orientational restraint between the principal vector of the decoupled POPC lipid and the normal axis of bilayer $\theta$ is applied so that the uncoupled lipid remains similar orientation as in fully coupled state. Thus, the volume sampled by uncoupled lipid in bulk bilayer is $2Z_RA(1 - Cos\theta)$ (3). $\Delta G_{FEP}^{bilayer}$ is the free energy contribution of decoupling one POPC lipids from a POPC bilayer. Similar as in the binding site, the flat-bottom harmonic restraints on $Z_R$ and $\theta$ have zero contribution when the lipid is fully coupled. Two different POPC bilayer sizes, 40 per leaflet and 60 per leaflet, were tested and yield consistent $\Delta G_{FEP}^{bilayer}$ values within uncertainty. In addition, $-k_B \mathrm{T} \ln(N_{lipid})$ is the free energy contribution due to the number of identical POPC lipids in the system, similar as the concept of the symmetric factor of a ligand.

**Table S3. The free energy contribution of each restraint for absolute binding energy calculation.**

| Free energy term | FEP (kcal/mol) | STD |
|---|---|---|
| $\Delta G_{FEP}^{site}$ | -160.9 | 0.1 |
| $\Delta G_{RMSD}^{gas}$ | 8.7 | 0.6 |
| $\Delta G_{geom}^{gas}$ | 2.3 | 0 |
| $\Delta G_{FEP}^{bilayer}$/lipid | -63.4 | 0.5 |
| $-k_B \mathrm{T} \ln(N_{lipid})$ | -3.4 | 0 |
| $\Delta G_{tot}$/lipid | (-160.9 + 8.7 + 2.3 + 3×63.4 )/3 -3.4 = 10.0 | 0.8 |

*The standard deviations for $e^{-\beta\Delta G_{FEP}^{site}}$ and $e^{+\beta\Delta G_{FEP}^{bilayer}}$ were calculated from MBAR. The number of identical POPC lipids in the system $N_{lipid}$ is 310. Restraint FEP was calculated using three independent simulations. $\Delta G_{geom}^{gas}$ is using the restraint constants regarding the bulk60 system.

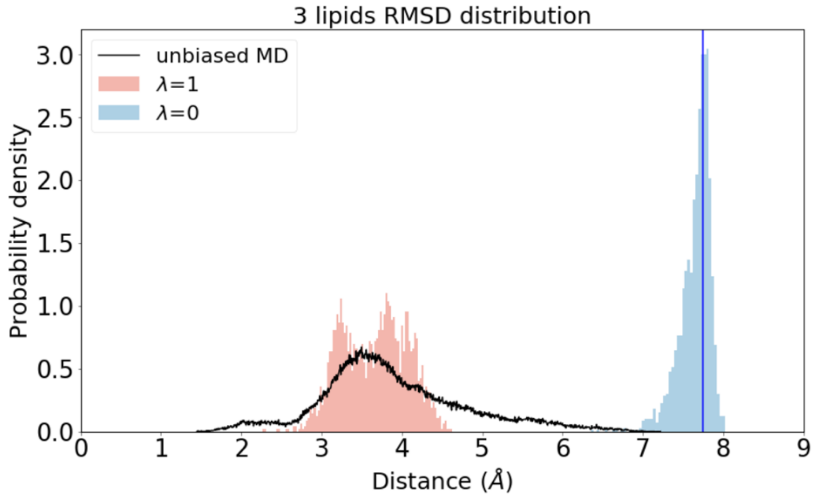


**Figure S1**. RMSD distributions of three lipids in the Piezo1 pore during FEP/ $\lambda$-REMD simulations ( $\lambda = 1$ and $\lambda = 0$) vs from unbiased MD simulations. The blue vertical line indicates the upper boundary of the flat-bottom RMSD harmonic restraint.

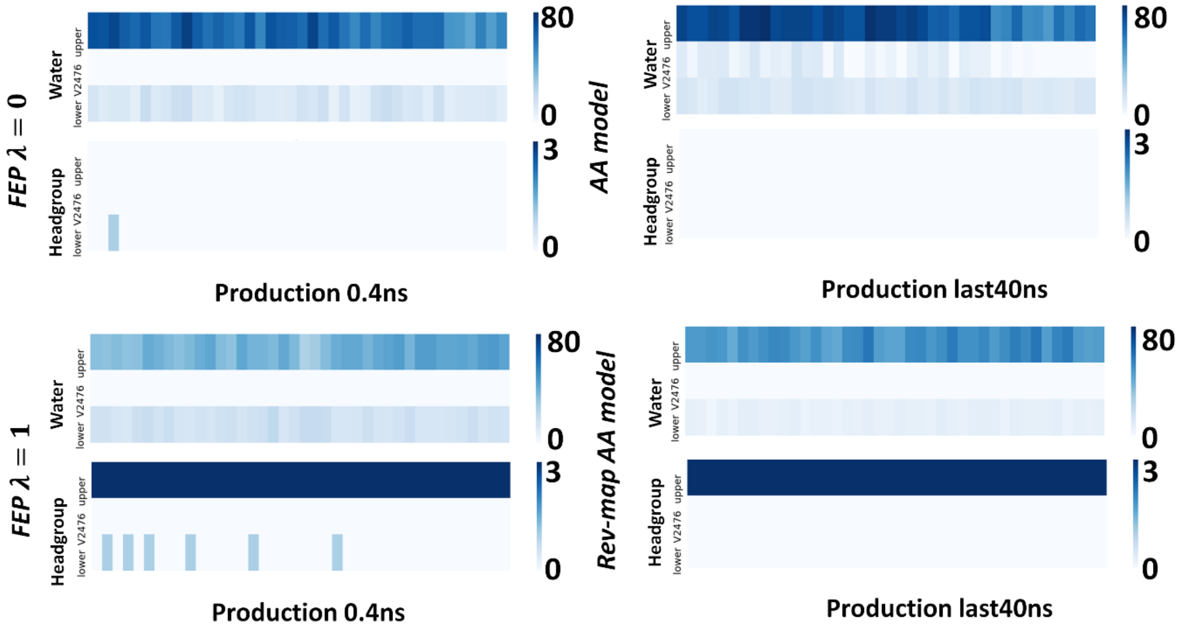


**Figure S2**. Water oxygen and headgroup phosphorus atom number distribution over time series in FEP/ $\lambda$-REMD simulations ( $\lambda = 1$ and $\lambda = 0$) vs. in AA model or CG Rev-mapped AA model.

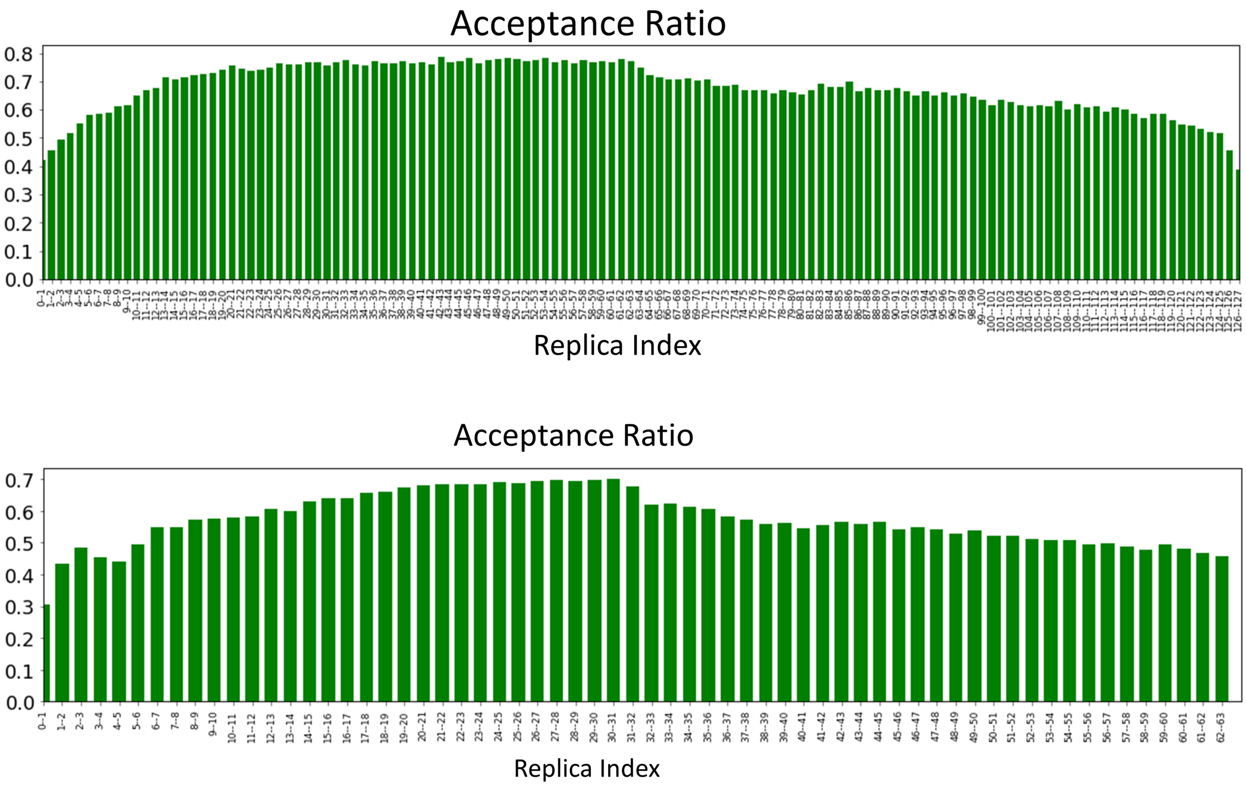


**Figure S3**. The acceptance ratio between each adject pair of 128 replicas in Piezo system (upper) and 64 replicas in bilayer-only during FEP/ $\lambda$-REMD simulations.

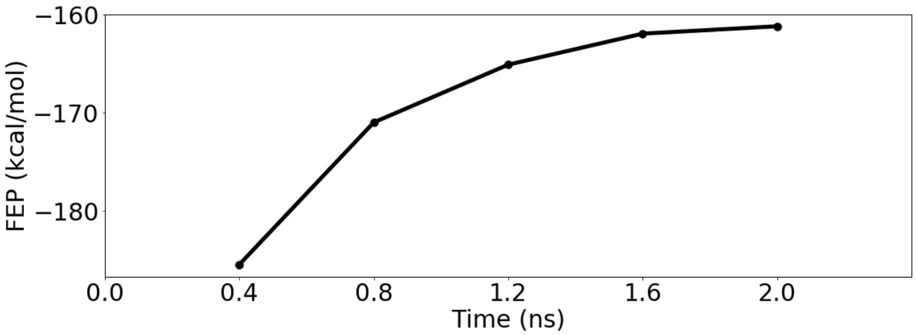

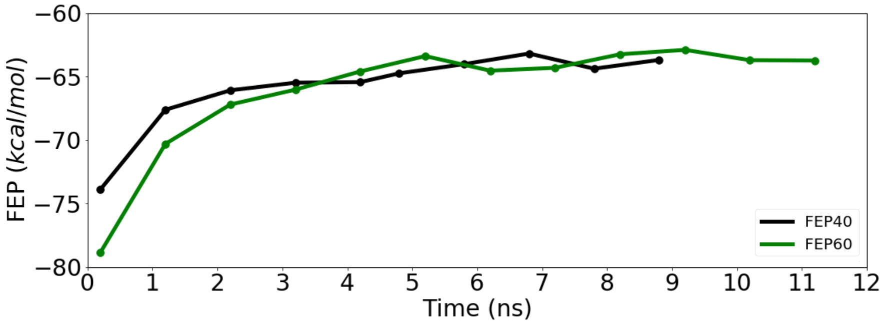


**Figure S4**. Convergence of the 3-lipid binding free energy calculations (upper) and bulk POPC bilayer size 40 and 60 systems (below) during FEP/ $\lambda$-REMD simulations.

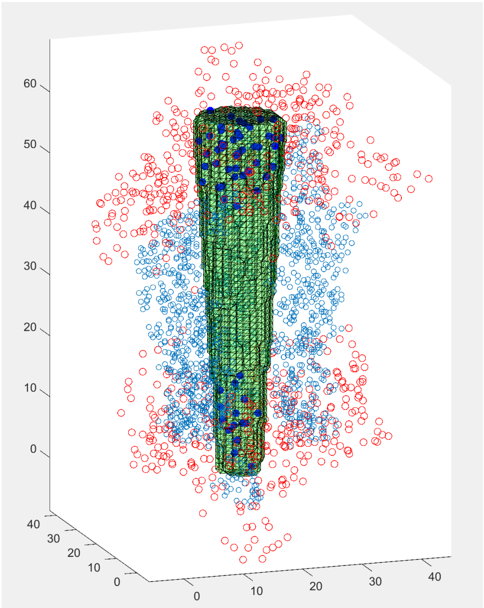

**Figure S5**. MATLAB classification plot for counting the water number in each pore region for $\lambda = 1$ during FEP/ $\lambda$-REMD simulations. Red dots stand for water oxygen atoms are within distance of 10Å from pore helix (residue P2455 to F2485, in cyan dots). Blue dots stand for the classified water oxygen atoms inside the cylinder pore region shown in grid green volume. Green frustum shape resembles the pore based on pore helix alpha carbon coordinates.

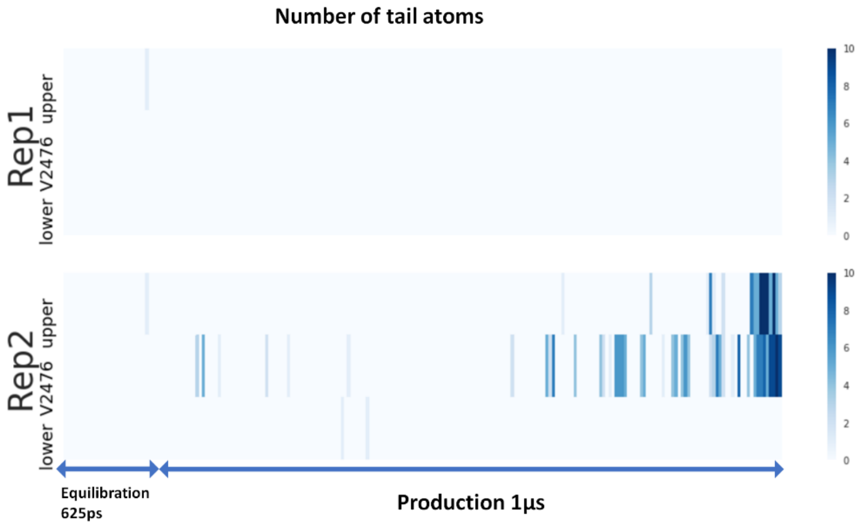


**Figure S6**. Tail atom number distribution for lower, hydrophobic constriction site and upper pore regions over time series in AA simulations.
